# Optimized Multicolour Immunofluorescence Panel for Cattle B Cell Phenotyping by an 8-Colour, 10-Parameter Panel

**DOI:** 10.1101/2021.10.25.465461

**Authors:** Eduard O. Roos, Marie Bonnet-Di Placido, William N Mwangi, Katy Moffat, Lindsay Fry, Ryan Waters, John A. Hammond

## Abstract

This 8-colour, 10-parameter panel has been optimised to distinguish between functionally distinct subsets of cattle B cells in both fresh and cryopreserved peripheral blood mononuclear cells (PBMCs). Existing characterised antibodies against cell surface molecules (immunoglobulin light chain (S-Ig(L)), CD20, CD21, CD40, CD71 and CD138) enabled the discrimination of 24 unique populations within the B cell population. This allows the identification of five putative functionally distinct B cell subsets critical to infection and vaccination responses; 1) naïve B cells (B_Naïve_), 2) regulatory B cells (B_Reg_), 3) memory B cells (B_Mem_), 4) plasmablasts (PB) and 5) plasma cells (PC). Although CD3 and CD8α can be included as an additional dump channel, it does not significantly improve the panel’s ability to separate “Classical” B cells. This panel will promote better characterisation and tracking of B cell responses in cattle as well as other bovid species as the reagents are likely to cross react.

## Background

As our knowledge of immune cell subsets and their functions increases, so does the need to identify and measure alterations in their phenotype and frequency. The mammalian B cell population consists of several functionally distinct subsets that together comprise the major mediator of humeral immunity (1,2). The development of naïve B cells (B_Naïve_) is important for long term immune protection (3–5). Driving the development of antibody secreting cells (ASC) and memory B cells (B_Mem_) is an essential requirement of many vaccines that elicit neutralizing antibody responses (6–8). Furthermore, these subsets are often the source of therapeutic antibody candidates (as vaccines or immunotherapies) against infectious diseases (6–8). Regulatory B cells (B_Reg_) also play a vital role in suppressing infectious diseases (9,10). Consequently, the identification and relative quantification of B cell subsets is a fundamental requirement when evaluating pathogen or vaccine induced immune responses and ultimately the development of better strategies to control diseases (1).

The capability to dissect B cell responses at high resolution is limited in many non-model species through a combination of limited reagents, lack of knowledge of species-specific B cell markers and standardised methods (11). This is certainly the case for cattle, a key food producing species and crucial for human nutrition globally, as a universal B cell lineage marker (i.e. CD19) and reagents against other well-known B cell subsets (e.g. IgD and CD38) are lacking (12). As technologies to design and deliver protective immunogens continue to emerge rapidly, it is essential to evaluate their applicability in other species as part of one health approaches. Consequently, a need to study cattle B cell responses and their maturation at a high resolution.

We have developed a flow cytometry panel using existing antibodies against cell surface markers based on knowledge in humans and mice (13). With no pan B cell markers known in cattle, such as CD19, CD72 or CD79α, we separated B cells from other lymphocytes using CD14 (CCG33, (14)) to exclude the monocytes, CD40 (IL-A158, (15)) as a B cell lineage marker, and included previously described cattle B cell markers such as CD21 (CC21, (16,17)) and surface immunoglobulin light-chain (S-Ig(L), IL-A58, (16,18)) (2,19). Subsets within these populations were further dissected using the activation and differentiation markers CD71 (IL-A165, (20)), CD20 (MEM-97, (21)) and CD138 (recombinant-F1.20/A, (Personal comm. Washington State University)).

Based on well characterised human and mouse B cell populations we hypothesise that these markers will identify five major subsets of B cells in cattle lymphocytes (Online Table 3): B_Naïve_, B_Mem_, B_Reg_, plasmablasts (PB) and plasma cells (PC) (13). The panel further allows for more in-depth characterisation of cattle B cells into 24 phenotypically unique subsets, following the gating strategy set out in Fig. 1; however, the functional discrimination and therefore importance between these subsets remains to be determined.

**Figure 1.**
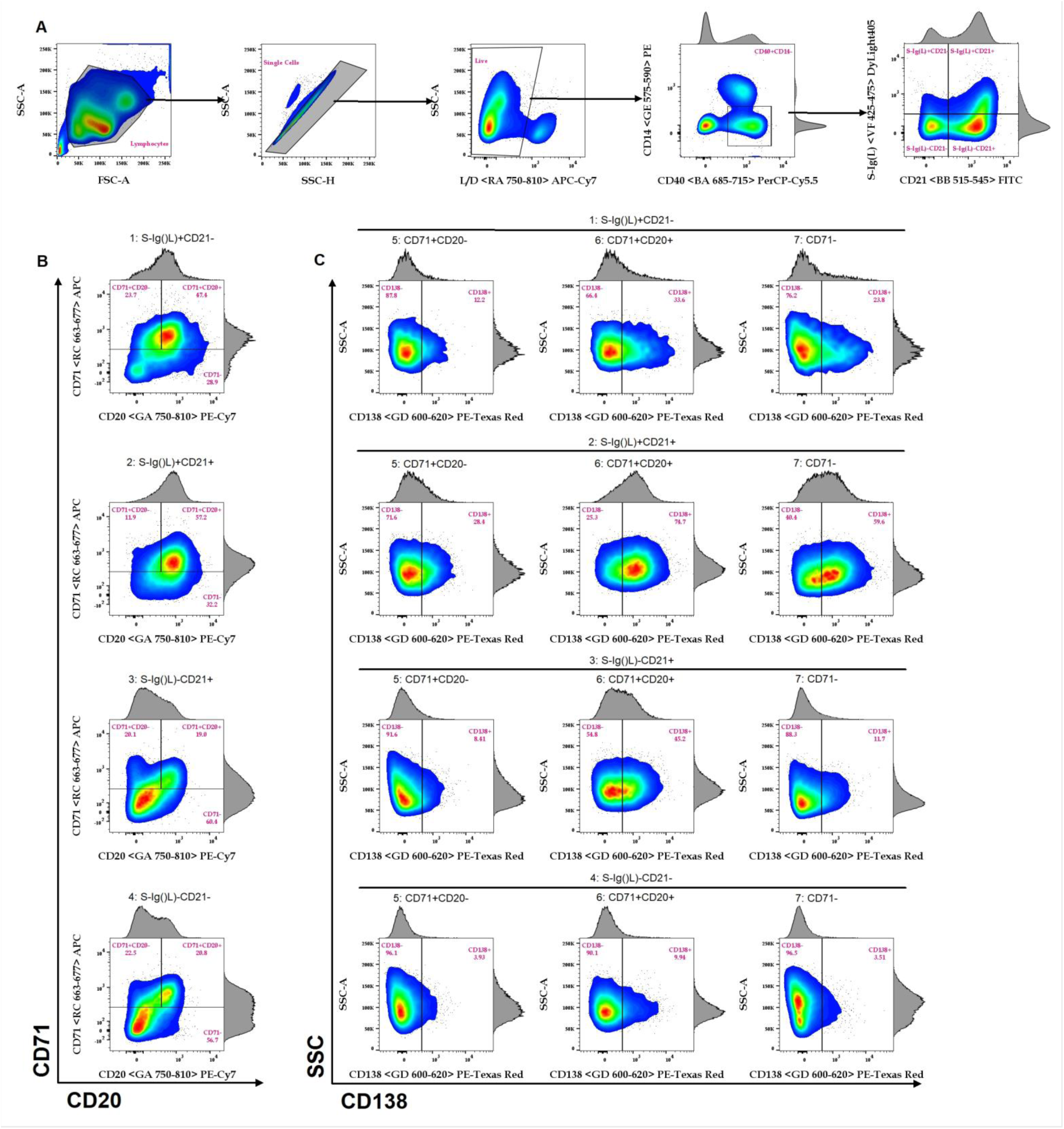
Gating strategy of the Cattle B cell panel into 24 minor subsets. A) The gating strategy followed from all events to the S-Ig(L) vs CD21 populations. B) S-Ig(L) vs CD21 further sub-gated into CD71 vs CD20 subsets for each of the four major subsets. C) The 24 minor subsets identified as the CD138^+^ and CD138^-^ from the three previous subsets in B. The parent population for each of the 24 minor subsets are listed above each gated population.

Our gating strategy consists of plotting CD40 against CD14 to select “classical” (CD40^+^CD14^-^) B cells. Although CD3 and CD8α are often used as a dump channel to isolate cattle B cells, their inclusion did not significantly improve separation (Online Figure 8). After identifying B cells, S-Ig(L) was plotted against CD21, allowing discrimination of four putative cattle B cell populations: CD21^-^S-Ig(L)^+^and CD21^+^S-Ig(L)^-^ single positive (SP), CD21^+^S-Ig(L)^+^ double positive (DP), and CD21^-^S-Ig(L)^-^ double negative (DN) cells (Fig. 1 A). Next, each population was further sub-divided by comparing CD71 against CD20 and sub-gated into CD71^+^ CD20^-^ SP, CD71^+^CD20^+^ DP and CD71^-^ populations (Fig.1 B). Lastly, each of these sub-gates were divided as either CD138^+^ or CD138^-^ (Fig.1 C), resulting in 24 minor subsets of cattle B cells. An important step while labelling the PBMCs is to first stain the cells with the CD20 antibody before adding any of the other antibodies in the panel (Online Figure 9).

Although this panel was developed and optimised on a BD LSRFortessa, it performed equally well using a BD Aria IIIU when sorting cattle B cell subsets for further molecular investigation. The panel also has potential to be adapted by moving the CD14 antibody into the Live/Dead channel (for example coupled to APC-Cy7), provided the monocytes do not need to be specifically analysed, or into an available violet channel, which will free the PE channel for an additional marker if needed (e.g. intra-cellular staining). The CD71 antibody in APC can also be moved into an available violet channel. Further potential to expand and tailor this panel includes the addition of a cattle cross-reactive human CD27 antibody, which could resolve the limitations in identifying B_Mem_ cells or the addition of an antibody against CD5, which is known to identify B1 cells in mice (22).

By identifying functional subsets of B cells this panel has the potential to dramatically improve our understanding of cattle immune responses to infection and vaccination, moving towards addressing some of the problems detailed in both Entrican *et al*. and Barroso *et al*., e.g. the lack of reagents to study the developmental cascade of cattle B cells (11,12). Additionally, the panel allows the enrichment or isolation of specific single B cells or their populations to further study function, specificity, and drive antibody discovery.

## Similarity to published OMIPs

None to date.

## Acknowledgements

The authors wish to acknowledge the valuable input of Dr. Kelcey Dinkel and Dr. Sally Madsen-Bouterse, as well as the exceptional technical support and animal care provided by Shelby Beckner, Megan Jacks, Emma Karel, Sarah Therrian, and Morgan Burke. This project has received funding from the European Union’s Horizon 2020 research and innovation programme under the VetBioNet grant agreement (731014). The authors would like to acknowledge the Flow Cytometry unit and the Immunological Toolbox unit supported by the United Kingdom Research and Innovation-Biotechnology and Biological Sciences Research Council awards (BBS/E/I/00007038 and BBS/E/I/00007039). Part of this work was supported by USAID (AID-BFS-P-13-00002) and the Bill and Melinda Gates Foundation (OPP1163607). The content of this manuscript is the sole responsibility of the USDA-ARS-ADRU and Washington State University and does not necessarily reflect the views of USAID or the United States Government.

---

**Table 1.**
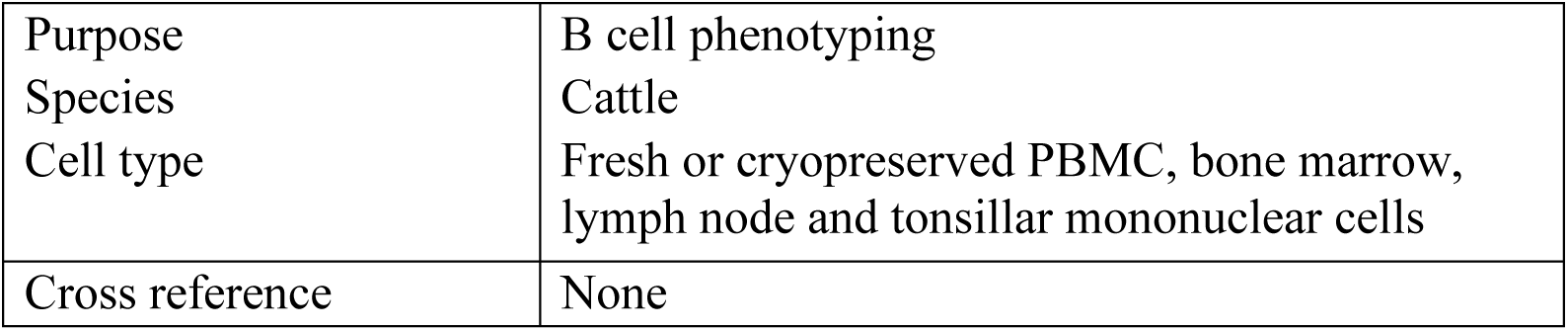
Summary table for Optimized Multicolour Immunofluorescence Panel

|  |  |
| --- | --- |
| Purpose | B cell phenotyping |
| Species | Cattle |
| Cell type | Fresh or cryopreserved PBMC, bone marrow, lymph node and tonsillar mononuclear cells |
| Cross reference | None |

**Table 2.**
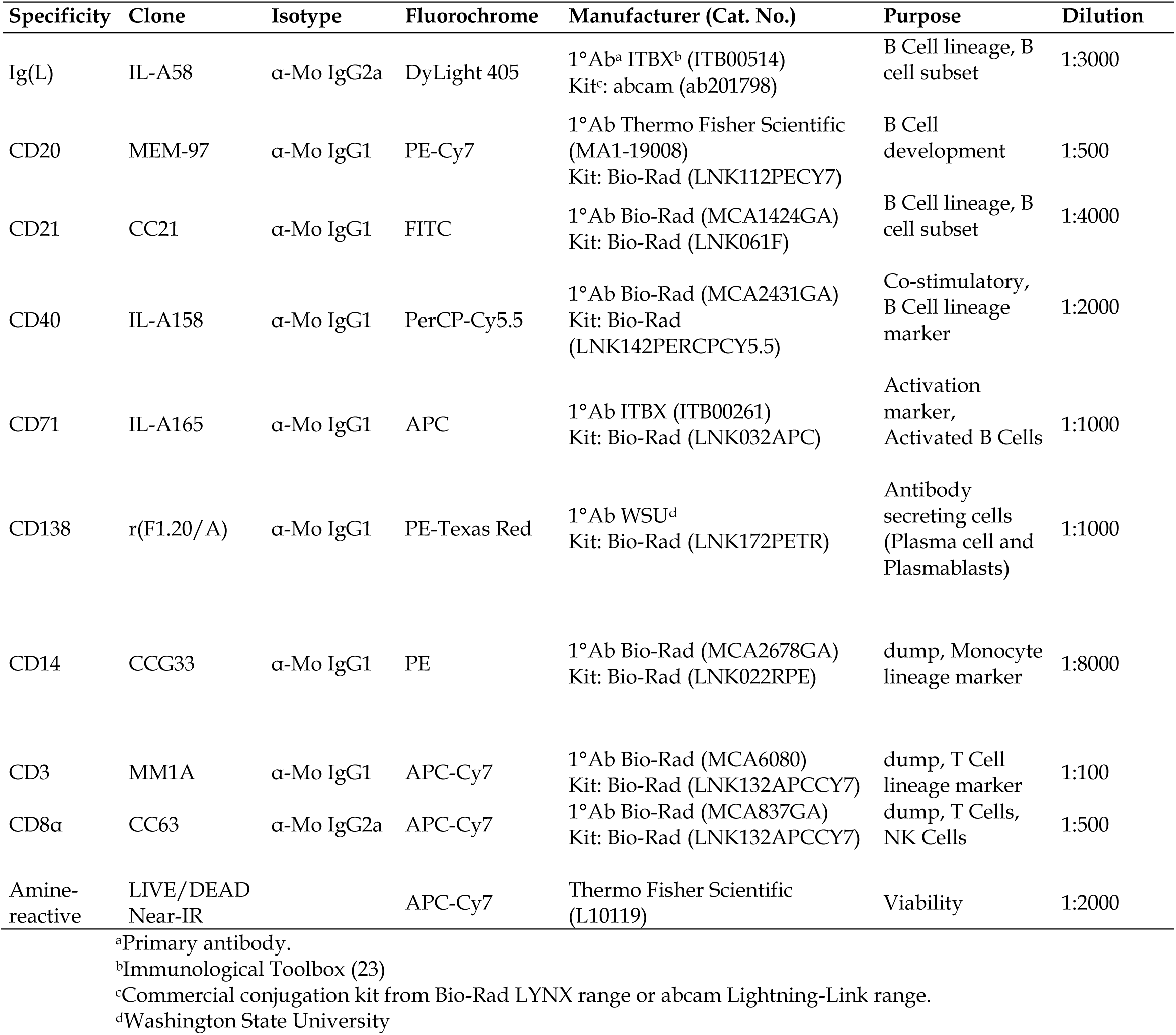
Reagents used for Optimized Multicolour Immunofluorescence Panel

## Literature Cited

1. Palm A-KE, Henry C. Remembrance of Things Past: Long-Term B Cell Memory After Infection and Vaccination. Front. Immunol. 2019;10:1–13.

2. LeBien TW, Tedder TF. B lymphocytes: how they develop and function. Blood 2008;112:1570–1580. Available at: https://ashpublications.org/blood/article/112/5/1570/25424/B-lymphocytes-how-they-develop-and-function.

3. Seifert M, Küppers R. Human memory B cells. Leukemia 2016;30:2283–2292. Available at: http://www.nature.com/articles/leu2016226.

4. Kaminski DA, Wei C, Qian Y, Rosenberg AF, Sanz I. Advances in Human B Cell Phenotypic Profiling. Front. Immunol. 2012;3:1–15. Available at: http://journal.frontiersin.org/article/10.3389/fimmu.2012.00302/abstract.

5. Nutt SL, Hodgkin PD, Tarlinton DM, Corcoran LM. The generation of antibody-secreting plasma cells. Nat. Rev. Immunol. 2015;15:160–171. Available at: http://dx.doi.org/10.1038/nri3795.

6. Lambour J, Naranjo-Gomez M, Piechaczyk M, Pelegrin M. Converting monoclonal antibody-based immunotherapies from passive to active: bringing immune complexes into play. Emerg. Microbes Infect. 2016;5:e92. Available at: http://dx.doi.org/10.1038/emi.2016.97.

7. Irani V, Guy AJ, Andrew D, Beeson JG, Ramsland PA, Richards JS. Molecular properties of human IgG subclasses and their implications for designing therapeutic monoclonal antibodies against infectious diseases. Mol. Immunol. 2015;67:171–182. Available at: http://dx.doi.org/10.1016/j.molimm.2015.03.255.

8. Bornholdt ZA, Turner HL, Murin CD, Li W, Sok D, Souders CA, Piper AE, Goff A, Shamblin JD, Wollen SE, Sprague TR, Fusco ML, Pommert KBJ, Cavacini LA, Smith HL, Klempner M, Reimann KA, Krauland E, Gerngross TU, Wittrup KD, Saphire EO, Burton DR, Glass PJ, Ward AB, Walker LM. Isolation of potent neutralizing antibodies from a survivor of the 2014 Ebola virus outbreak. Science (80-.). 2016;351:1078–1083. Available at: https://www.science.org/doi/10.1126/science.aad5788.

9. Shen P, Fillatreau S. Suppressive functions of B cells in infectious diseases: Fig. 1. Int. Immunol. 2015;27:513–519. Available at: https://academic.oup.com/intimm/article-lookup/doi/10.1093/intimm/dxv037.

10. Rosser EC, Mauri C. Regulatory B Cells: Origin, Phenotype, and Function. Immunity 2015;42:607–612. Available at: http://dx.doi.org/10.1016/j.immuni.2015.04.005.

11. Entrican G, Lunney JK, Wattegedera SR, Mwangi W, Hope JC, Hammond JA. The Veterinary Immunological Toolbox: Past, Present, and Future. Front. Immunol. 2020;11:1–8. Available at: https://www.frontiersin.org/article/10.3389/fimmu.2020.01651/full.

12. Barroso R, Morrison WI, Morrison LJ. Molecular Dissection of the Antibody Response: Opportunities and Needs for Application in Cattle. Front. Immunol. 2020;11:1–10. Available at: https://www.frontiersin.org/article/10.3389/fimmu.2020.01175/full.

13. Murphy K, Weaver C. Janeway’s Immunobiology. 9th ed. (Murphy K, Weaver C, editors). New York, NY: Garland Science/Taylor & Francis; 2018. 1–904 p. Available at: https://www.journals.uchicago.edu/doi/10.1086/696793.

14. Sopp P, Kwong LS, Howard CJ. Identification of bovine CD14. Vet. Immunol. Immunopathol. 1996;52:323–328. Available at: https://linkinghub.elsevier.com/retrieve/pii/0165242796055833.

15. Glew EJ, Carr B V., Brackenbury LS, Hope JC, Charleston B, Howard CJ. Differential effects of bovine viral diarrhoea virus on monocytes and dendritic cells. J. Gen. Virol. 2003;84:1771–1780. Available at: https://www.microbiologyresearch.org/content/journal/jgv/10.1099/vir.0.18964-0.

16. Naessens J, Newson J, McHugh N, Howard CJ, Parsons K, Jones B. Characterization of a bovine leucocyte differentiation antigen of 145,000 MW restricted to B lymphocytes. Immunology 1990;69:525–30. Available at: http://www.ncbi.nlm.nih.gov/pubmed/2185984.

17. Naessens J, Hopkins J. Introduction and summary of workshop findings. Vet. Immunol. Immunopathol. 1996;52:213–235. Available at: https://linkinghub.elsevier.com/retrieve/pii/0165242796055663.

18. Williams DJL, Newson J, Naessens J. Quantitation of bovine immunoglobulin isotypes and allotypes using monoclonal antibodies. Vet. Immunol. Immunopathol. 1990;24:267–283. Available at: https://linkinghub.elsevier.com/retrieve/pii/016524279090042Q.

19. Cyster JG, Allen CDC. B Cell Responses: Cell Interaction Dynamics and Decisions. Cell 2019;177:524–540. Available at: https://doi.org/10.1016/j.cell.2019.03.016.

20. Naessens J, Davis WC. Ruminant cluster CD71. Vet. Immunol. Immunopathol. 1996;52:257–258. Available at: https://linkinghub.elsevier.com/retrieve/pii/0165242796055729.

21. Faldyna M, Samankova P, Leva L, Cerny J, Oujezdska J, Rehakova Z, Sinkora J. Cross-reactive anti-human monoclonal antibodies as a tool for B-cell identification in dogs and pigs. Vet. Immunol. Immunopathol. 2007;119:56–62. Available at: https://linkinghub.elsevier.com/retrieve/pii/S0165242707002152.

22. Cossarizza A, Chang H-D, Radbruch A, Acs A, Adam D, Adam Klages S, Agace WW, Aghaeepour N, Akdis M, Allez M, Almeida LN, Alvisi G, Anderson G, Andrä I, Annunziato F, Anselmo A, Bacher P, Baldari CT, Bari S, Barnaba V, Barros Martins J, Battistini L, Bauer W, Baumgart S, Baumgarth N, Baumjohann D, Baying B, Bebawy M, Becher B, Beisker W, Benes V, Beyaert R, Blanco A, Boardman DA, Bogdan C, Borger JG, Borsellino G, Boulais PE, Bradford JA, Brenner D, Brinkman RR, Brooks AES, Busch DH, Büscher M, Bushnell TP, Calzetti F, Cameron G, Cammarata I, Cao X, Cardell SL, Casola S, Cassatella MA, Cavani A, Celada A, Chatenoud L, Chattopadhyay PK, Chow S, Christakou E, ČiČinŠain L, Clerici M, Colombo FS, Cook L, Cooke A, Cooper AM, Corbett AJ, Cosma A, Cosmi L, Coulie PG, Cumano A, Cvetkovic L, Dang VD, Dang Heine C, Davey MS, Davies D, De Biasi S, Del Zotto G, Dela Cruz GV, Delacher M, Della Bella S, Dellabona P, Deniz G, Dessing M, Di Santo JP, Diefenbach A, Dieli F, Dolf A, Dörner T, Dress RJ, Dudziak D, Dustin M, Dutertre C, Ebner F, Eckle SBG, Edinger M, Eede P, Ehrhardt GRA, Eich M, Engel P, et al. Guidelines for the use of flow cytometry and cell sorting in immunological studies (second edition). Eur. J. Immunol. 2019;49:1457–1973. Available at: http://doi.wiley.com/10.1002/eji.201646632.

23. Mwangi W, Maccari G, Hope JC, Entrican G, Hammond JA. The UK Veterinary Immunological Toolbox Website: promoting vaccine research by facilitating communication and removing reagent barriers. Immunology 2020;161:25–27. Available at: http://doi.wiley.com/10.1111/imm.13227.

