## Supplementary material for "Optimized Multicolour Immunofluorescence Panel for Cattle B Cell Phenotyping by an 8-Colour, 10-Parameter Panel": Online supporting information

**Online Table 1. Instrument configuration.**

| Laser Wavelength (nm) | Laser Power (mW) | Laser Type | Detector | Spectral Range for Detector (nm) | Dichroic LP Filter (nm) | Band Pass (nm) | Example Fluorochrome |
|---|---|---|---|---|---|---|---|
| 640 (Red – R) | 40 | COHERENT CUBE 640-40C | A | 750-810 | 750 | 780/60 | Live/Dead NIR |
| | | | B | 710-753 | 710 | 730/45 | AF700 |
| | | | C | 663-677 | --- | 670/14 | APC |
| 561 (Green – G) | 50 | COHERENT Sapphire 561 LP | A | 750-810 | 750 | 780/60 | PE-Cy7 |
| | | | B | 685-735 | 685 | 710/50 | PE-Cy5.5 |
| | | | C | 655-685 | 635 | 670/30 | PE-Cy5 |
| | | | D | 600-620 | 600 | 610/20 | PE-Texas Red |
| | | | E | 575-590 | --- | 582/15 | PE |
| 488 (Blue – B) | 50 | COHERENT Sapphire 488 LP | A | 685-715 | 685 | 695/40 | PerCP-Cy5.5 |
| | | | B | 515-545 | 505 | 530/30 | FITC |
| 405 (Violet – V) | 50 | COHERENT CUBE 405-50C | A | 750-810 | 750 | 780/60 | BV786 |
| | | | B | 690-735 | 690 | 710/50 | BV711/SB700 |
| | | | C | 650-670 | 635 | 660/20 | BV650 |
| | | | D | 595-620 | 595 | 610/20 | BV605/SB600 |
| | | | E | 500-550 | 475 | 525/50 | ER-T White/Blue |
| | | | F | 425-475 | --- | 450/50 | BV421/DyLight405 |

The Optimized Multicolor Immunofluorescence Panel was optimized on a LSRFortessa cytometer (BD Biosciences). The cytometer had a four-laser configuration, with the listed optical configuration.

**Online Table 2.** Different versions of the panel as it underwent changes in antibody and fluorochrome combinations.

| Antigen | Clone | Isotype | Version 1 | Version 2 | Version 3 | Version 4 | Version 5 | Version 6 | Version 7 |
|---|---|---|---|---|---|---|---|---|---|
| Ig(L) | IL-A58 | IgG2a | DyLight405 | DyLight405 | DyLight405 | DyLight405 | DyLight405 | DyLight405 | DyLight405 |
| CD20 | MEM-97 | IgG1 | PE-Cy5 | PE-Cy5 | PE-Cy5 | PE-Cy5 | PE | PE/PE-Cy5 | PE-Cy7 |
| CD21 | CC21 | IgG1 | PE | FITC | FITC | FITC | PE-Cy5 | FITC | FITC |
| CD14 | CCG33 | IgG1 | FITC | APC | PerCP | PE | PE-Cy7 | PE-Cy7 | PE |
| CD40 | IL-A158 | IgG1 | PE-Cy7 | PE | PE | PerCP | PerCP-Cy5.5 | PerCP-Cy5.5 | PerCP-Cy5.5 |
| CD71 | IL-A165 | IgG1 | APC | PE-Cy7 | PE-Cy7 | PE-Cy7 | APC | APC | APC |
| CD3<br>CD8α | MM1A<br>CC63 | IgG1<br>IgG2a | FITC | APC | APC | APC | FITC | APC-Cy7 | APC-Cy7 |
| L/D NIR | N/A | N/A | APC-Cy7 | APC-Cy7 | APC-Cy7 | APC-Cy7 | APC-Cy7 | APC-Cy7 | APC-Cy7 |
| CD138 | F1.20/A | rAb IgG1 | N/A | N/A | PE-Tex Red | PE-Tex Red | PE-Tex Red | PE-Tex Red | PE-Tex Red |

Summary of changes made to optimise the panel:
Version 1: Original panel design.
Version 2: Moved dump to APC (included CD14); CD71 to PE-Cy7; CD40 to PE; CD21 to FITC.
Version 3: Addition of CD138 on PE-Texas Red; Changed CD14 to PerCP from APC dump.
Version 4: Swoped CD14 to PE and CD40 to PerCP.
Version 5: Moved dump to FITC; CD71 to APC; CD14 to PE-Cy7; CD20 to PE; CD21 to PE-Cy5; CD40 to PerCP-Cy5.5.
Version 6: Moved dump to APC-Cy7 (Live/Dead); CD21 to FITC, additional test to confirm the exclusion of PE-Cy5 was done using CD20 on PE-Cy5.
Version 7: Swoped CD20 to PE-Cy7 and CD14 to PE.

**Online Table 3.** Five potential cattle B cell subsets of major importance.

| **Subset** | **Phenotype** | **References** |
|---|---|---|
| Naïve B Cells | $CD40^{+}CD14^{-}CCD21^{+}CD20^{low/-}CD71^{low/-}CD138^{-}$ | (1) |
| Memory B Cells | $CD40^{+}CD14^{-}$ $CD21^{+}S\text{-}Ig(L)^{+}CD20^{+}CD71^{+}CD138^{-}$ | (1–4) |
| Regulatory B Cells | $CD40^{+}CD14^{-}CD21^{+}S\text{-}Ig(L)^{low/+}CD20^{+}CD71^{+}CD138^{+}$ | (1,2) |
| Plasmablasts | $CD40^{+}CD14^{-}CD21^{low/+}S\text{-}Ig(L)^{low/-}CD20^{-}CD71^{+}CD138^{-}$ | (1–3) |
| Plasma Cells | $CD40^{+}CD14^{-}CD21^{low/+}S\text{-}Ig(L)^{low/-}CD20^{-}CD71^{+}CD138^{+}$ | (1–3) |

**Online Table 4.** Antibodies tested but not used for the final panel.

| Specificity | Fluorochrome[1] | Ab clone | Vendor | Dilution | Reason for exclusion |
|---|---|---|---|---|---|
| CD71 | AF488 | IL-A77 | ITBX | 1:10 | Not available anymore. |
| S-Ig(L) | AF488 | IL-A59 | ITBX | 1:1000 | Comparable staining profile to IL-A58. |
| CD21 | AF488 | CC51 | ITBX | 1:1000 | Comparable staining profile to CC21. |
| CD14 | AF488 | CAM36A | Bio-Rad | 1:1000 | Same staining profile as CCG33 which is available through the Immunological Toolbox. |

[1] Alexa Fluor 488 was a secondary whole IgG antibody used at 1:2000 for initial screening.

**Developmental Strategy**

This eight-colour B-cell panel was designed for a BD Fortessa LSR with the configuration as outlined in the Online Table 1. We prioritized mouse anti-cattle monoclonal antibodies (mAb) that were also reactive in other ruminant species to expand the potential utility in the future with minimal changes required. Based on data from other species, this panel will identify basic cattle B-cell subsets such a naïve ($M_{Naïve}$), memory ($B_{Mem}$), regulatory ($B_{Reg}$) and antibody secreting cells (ASC; Plasmablasts (PB) and Plasma cells (PC)) (Online Table 3). The panel also identifies additional subsets of B cells that need to be further investigated and validated with regards to their functionality (Fig. 1).

Since most of the mAbs are not commercially available in the desired fluorochromes, this panel was further limited to fluorochromes that can be conjugated in house (we refrained from custom conjugations to ensure accessibility to other laboratories). The in-house conjugation kits used were the LYNX range (Bio-Rad) as well as the Lightning-Link range (abcam) listed in Table 2. These restrictions further complicated the panel design, as some of these cattle markers are dim (CD20, CD71, CD3) or smear (S-Ig(L), CD20, CD71) populations and required several optimisation steps, i.e. conjugated to various fluorochromes and tested in the panel (Online Table 2). The limited number of cattle reagents made it difficult to design and optimise the panel as only a single cattle CD3 antibody and a single cattle CD71 antibody clone is available, the recombinant cattle CD138 antibody that we used is reported for the first time and the human CD20 antibody was the only CD20 mAb that is crossreactive in cattle (no cattle CD20 antibody is available). The availability of selected mAbs was based on their listing on the UK Immunological Toolbox website as commercially available or available to request (5).

*Version 1 and 2*

The initial intention for the panel was to include CD3, CD8α and CD14 in a single dump channel in FITC (BB 515-545 detector, Version 1) or APC (RC 663-677 detector, Version 2) (Online Table 2). The separation of CD40 was good in both versions and especially in version 2 with the dump channel on APC (RC 663-677 detector, Online Figure 1). While the separation

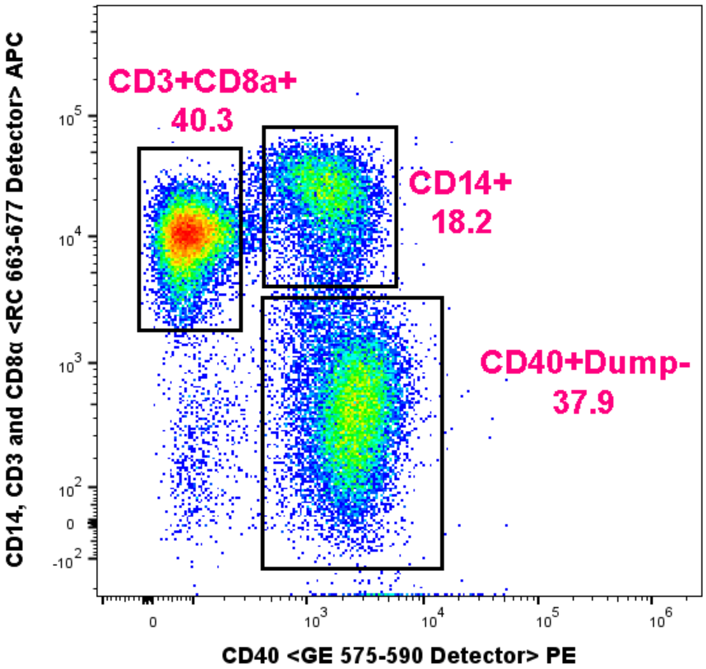


**Online Figure 1**
Isolation of CD40⁺ (PE) from all live events against the dump channel (CD14, CD3 and CD8α on APC; version 2).

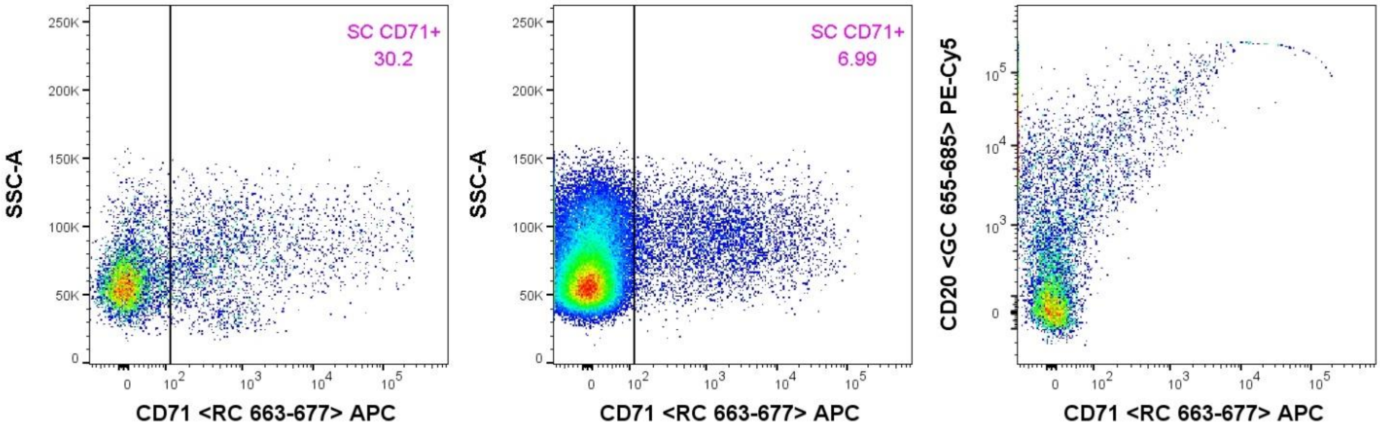


**Online Figure 2**
Spectral spread of CD20 (PE-Cy5) into CD71 (APC) reduces the discrimination of the CD71⁺ populations after applying compensation. Left: Single colour stain of CD71 (APC); Middle: Full panel stain of version 1 after applying compensation, indicating the reduce resolution of CD71 (APC); Right: Single colour stain of CD20 (PE-Cy5) indicating the spectral spread into the APC.

of the CD40 and dump channel was acceptable in version 1, the CD71 (APC, RC 663-677 detector) and CD20 (PE-Cy5, GC 655-685 detector) had high spectral spread, with most of the spectral spread of PE-Cy5 going into the APC (RC 663-677 detector) and reducing the resolution of the CD71⁺ population in the fully stained sample (Online Figure 2). Therefore, the panel was changed to improve the resolution of the dump channel and that of the CD71⁺ population.

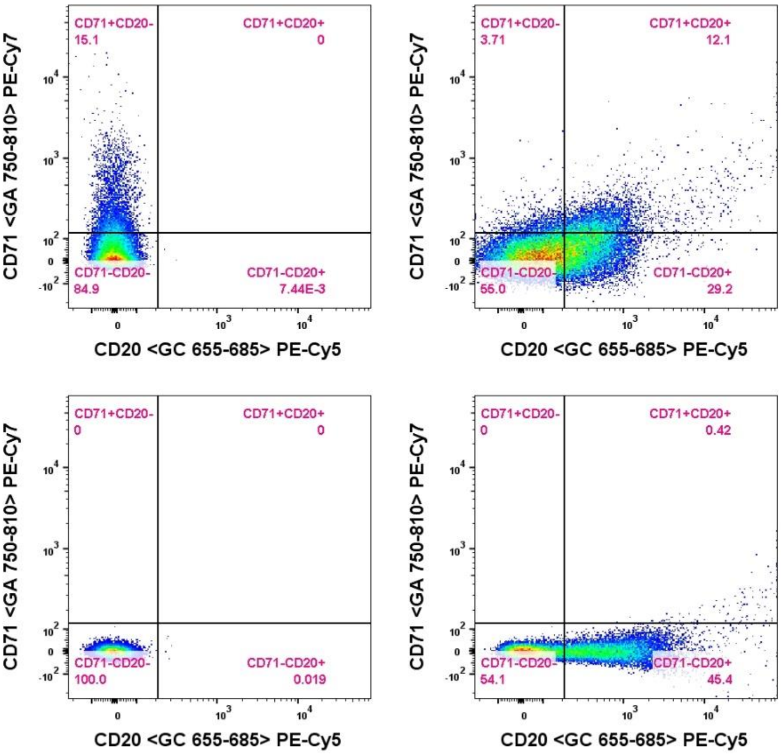


**Online Figure 3**
Representative sample of the spectral spread reduing the resolution of CD71 (PE-Cy7) and CD20 (PE-Cy5) after applying compensation. Top left: Single colour stain of CD71 (PE-Cy7); Top right: Full panel stain of version 2 after applying compensation, indicating the reduce resolution of CD71 and CD20; Bottom left: unstained sample; Bottom right: Single colour stain of CD20 (PE-Cy5).

These changes required a complete reshuffle of the antibody and fluorochrome combinations (Online Table 2). Thus, in version 2 the dump channel was moved to APC (RC 663-677 detector) to reduce the impact of the spectral spread into APC (RC 663-677 detector). The CD40 antibody was moved from PE-Cy7 (GA 750-810 detector) to PE (GE 575-590 detector), ensuring the maximum separation between the dump channel and CD40. The CD21 antibody was moved from PE (GE 575-590 detector) to FITC (BB 515-545 detector) and CD71 antibody was changed from APC (RC 663-677 detector) to PE-Cy7 (GA 750-810 detector). Despite good separation of CD40 and the dump channel, there were further concerns with CD20 (PE-Cy5, GC 655-685 detector) and CD71 (PE-Cy7, GA 750-810 detector). The CD71+ and CD20+ populations could not be fully resolved in version 2 to 4 of the panel (Online Figure 3).

As the panel had to be changed again, the CD14 antibody was moved into its own channel from version 3 onwards to allow the study of monocytes if desired. However, in this panel, CD14 is primarily included for the separation of B cells from monocytes. Therefore, it could be added to the LIVE/DEAD NIR (RA 750-810 detector) if desired (discussed further under Hints and Tips). The dump channel only included CD3 and CD8α from this version onward.

*Version 3 and 4*

Both CD14 (Version 3) and CD40 (Version 4) antibodies were tested on PerCP (BA 685-715 detector). The PerCP (BA 685-715 detector) was too dim to resolve these populations (Online Figure 4). Therefore, we changed the CD40 antibody to PerCP-Cy5.5 (BA 685-715 detector). This resolved the separation of the $CD40^+$ population and it was kept on this fluorochrome until the final panel (Online Table 2). Additionally, and CD138 antibody was also included in the panel from version 3 onward and kept on PE-Texas Red (GD 600-620 detector) as an additional differentiation marker until the final version.

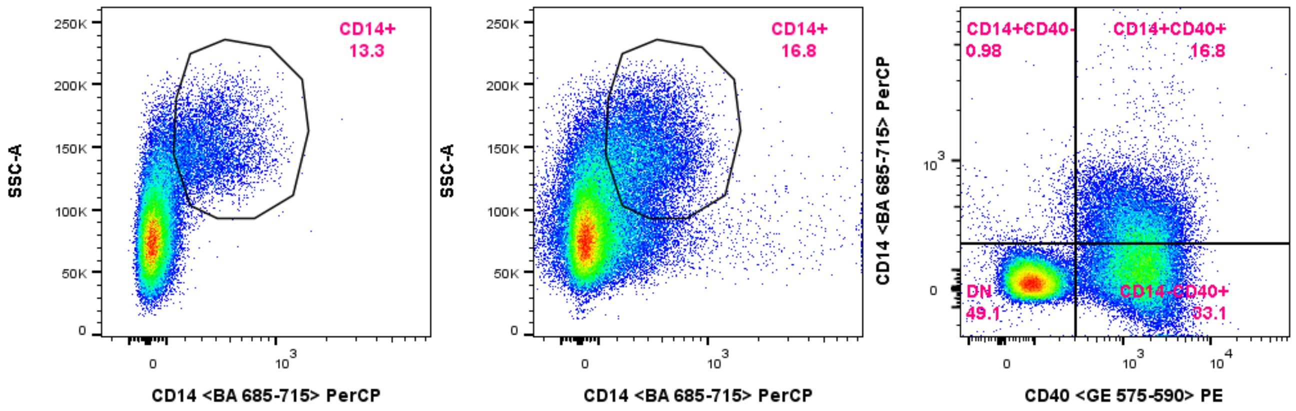


**Online Figure 4**
Version 3 as representative of how dim the PerCP (BA 685-715) was to differentiate the CD14 (CD40 not shown). Left: CD14 (PerCP) single colour. Middle: All antibodies for version 3 and gating for the CD14$^{+}$ population. Right: Resolution between CD14 (PerCP) and CD40 (PE).

*Version 5*

The panel was changed from version 4 in an attempt to improve the poor resolution of CD71 (PE-Cy7, GA 750-810 detector) and CD20 (PE-Cy5, GC 655-685 detector) populations (Online Figure 3). These two antibodies were placed on APC (RC 663-677 detector) and PE (GE 575-590 detector), respectively. This caused a cascade of changes in version 5 of the panel (Online Table 2). However, the spectral spread from CD21, now on PE-Cy5, caused difficulties in resolving CD71 in APC (RC 663-677 detector), as discussed below.

The initial testing of the CD40 antibody on PerCP-Cy5.5 (BA 685-715 detector) was promising but further investigation determined that the spectral spread from the CD21 antibody on PE-Cy5 was too high into the PerCP-Cy5.5 (BA 685-715 detector), PE-Cy7 (GA 750-810 detector) and APC (RC 663-677 detector) channels (Online Figure 5). The spectral spread from PE-Cy5 into other channels reduced the ability to properly resolve these subsets. Thus, version 5 of the panel (Online Table 2) was

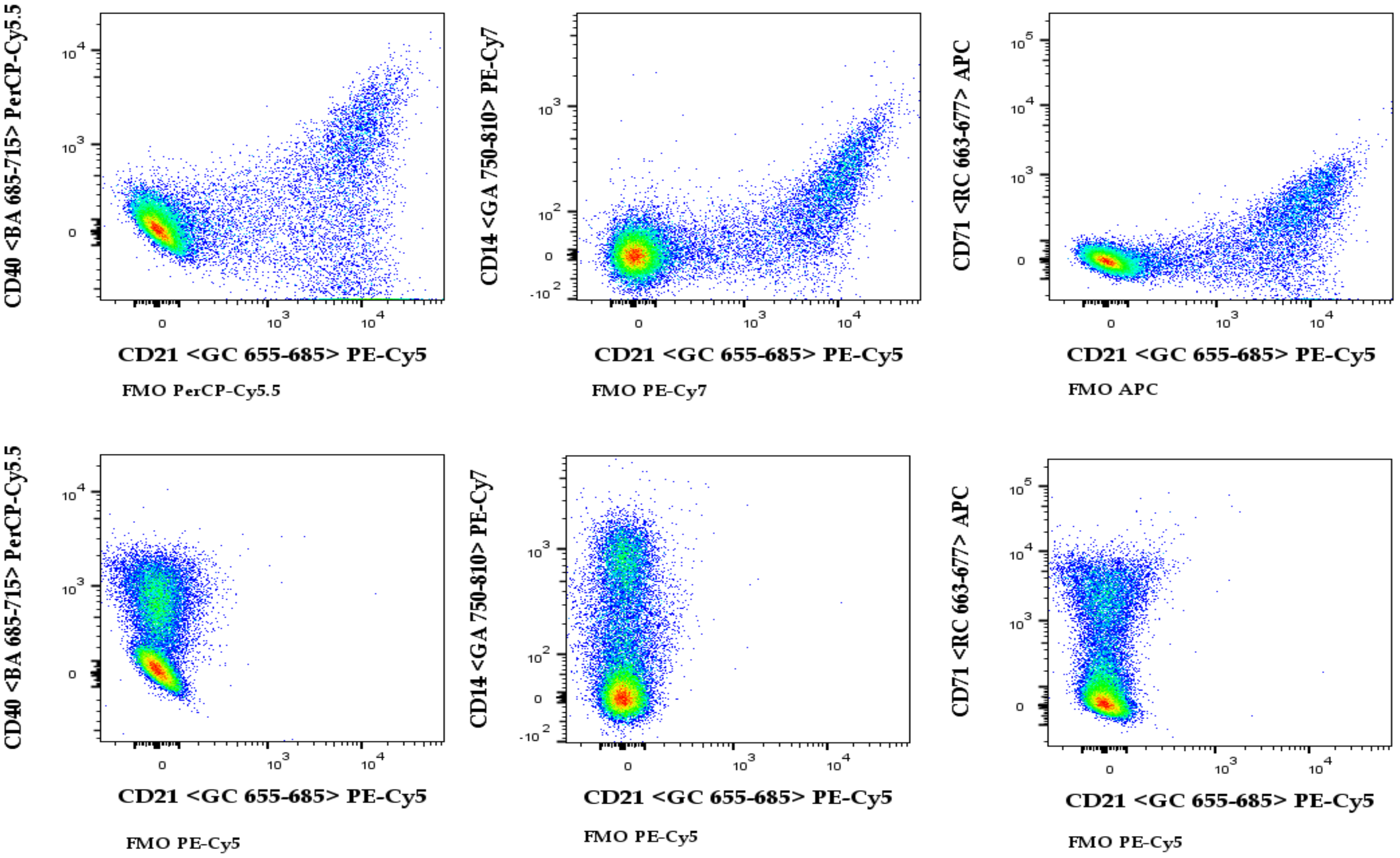


**Online Figure 5** Spectral spread observed of PE-Cy5 after compensation into PerCP-Cy5.5, PE-Cy7 and APC comparing the FMO of each of the fluorochromes against PE-Cy5.

changed since it allowed us to have our most difficult-to-resolve populations on separate laser lines. This reduced the spectral spread experienced with PE-Cy5 specifically. This change reduced the compensation to be applied for the following versions of the panel (Online Table 2). Additionally, the PE-Cy5 spectral spread into PerCP-Cy5.5 (BA 685-715 detector) and PE-Cy7 (GA 750-810 detector) decreased the discriminatory power between CD40 and CD14 (Online Figure 6 A and B), reducing the ability to distinguish B cells from monocytes. This discrimination was improved in the final versions of the panel by completely removing PE-Cy5 (GC 655-685 detector) from our panel (Online Figure 6 C, Online Table 2). The change of the CD21 antibody from PE-Cy5 (GC 655-685 detector) required the dump channel to be changed to the LIVE/DEAD NIR (APC-Cy7, RA 750-810 detector) to open up FITC (BB 515-545 detector) for the CD21 antibody in the panel (Online Table 2).

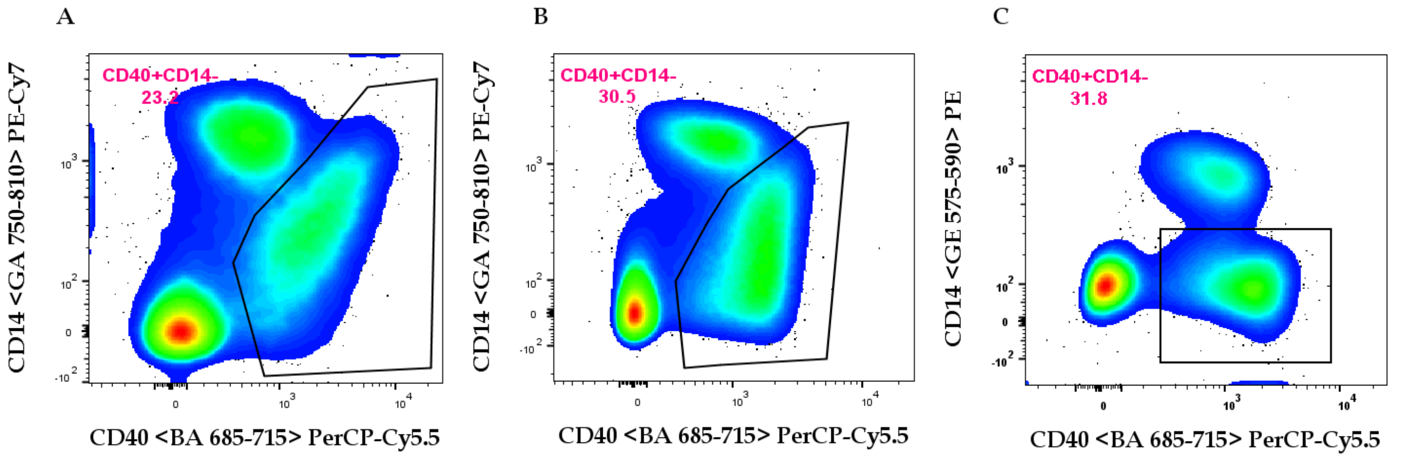


**Online Figure 6** Spectral spread observed of PE-Cy5 after compensation and its influence on the discriminatory power between CD40 and CD14 with CD21 (A, version 5) or CD20 (B, version 6 additional test) antibodies on PE-Cy5. The final version for the panel had no PE-Cy5 (C, version 7) and the CD14 antibody on PE.

*Version 6 and 7*

The use of CD20 and CD138 antibodies on PE (GE 575-590 detector) and PE-Texas Red (GD 600-620 detector) respectively, showed some spectral spread from PE into the PE-Texas Red (GD 600-620 detector, Online Figure 7 A). The loss of the CD20 signal was further investigated as discussed later. Consequently, we tried to avoid markers that are highly expressed on B cells in the PE channel.

As a result, the CD14 antibody was conjugated to PE (GE 575-590 detector) to determine what the impact of the PE spectral spread into PE-Texas Red (GD 600-620 detector) would be (Online Figure 7 B). And so, the CD20 antibody was swapped out with the CD14 antibody that was on PE-Cy7 (GA 750-810 detector) (Online Table 2).

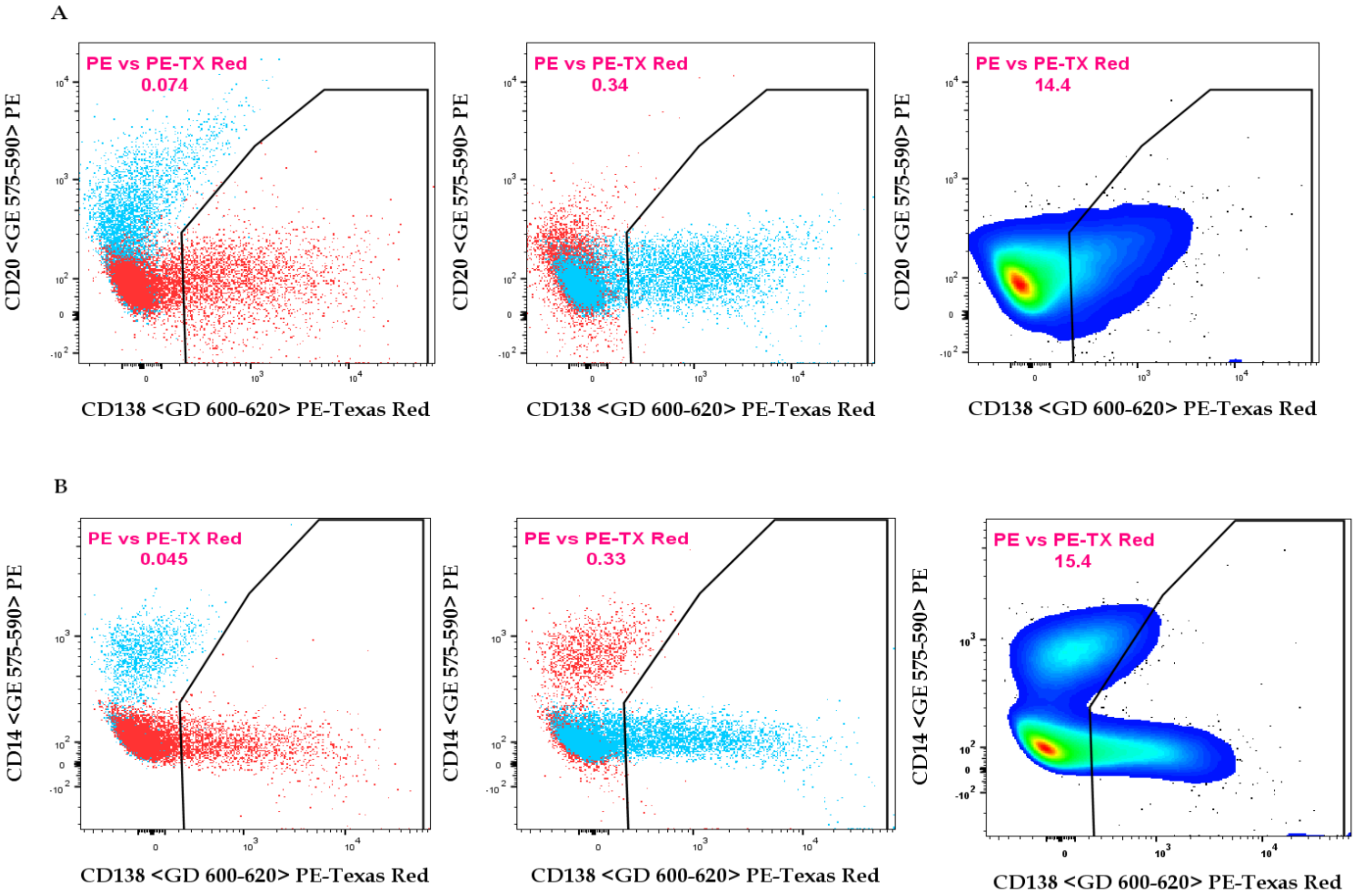


**Online Figure 7** Spectral spread observed of PE against the PE-Texas Red. A) Version 6 spectral spread from the of PE (CD20) into PE-Texas Red (CD138). B) Version 7 spectral spread from the of PE (CD14) into PE-Texas Red (CD138). Red represents the FMO for each of the fluorochrome and the light blue represents the single colour stain. From left to right; PE, PE-Texas Red and full stain, respectively.

In the end, the CD14 antibody on PE (GE 575-590 detector) was selected as it is not present on B cells, and thus allowed for better resolution in the panel to discriminate B cells and monocytes (Online Figure 6 C). The dump channel was moved to APC-Cy7 (same as LIVE/DEAD NIR) from version 6 onwards, but as discussed below, its inclusion did not make a significant difference in distinguishing the B cells

(CD40$^+$CD14$^-$). Spectral spread was the main reason antibodies were changed to different fluorochromes in this panel.

*Hints and tips*

The inclusion of an CD40, a co-stimulatory molecule, was used to help define the B cell population. Given that there is no definitive B cell marker in cattle, CD40 was used to allow the separation of the dump channel (CD3$^+$CD8α$^+$), monocytes (CD14$^+$) and B cells (CD40$^+$CD14$^-$). However, we decided to leave the dump channel out of the final analysis of the panel since it made no significant difference with regard to the CD40$^+$ population (Online Figure 8), and consequently on the isolation of B cells. Nonetheless, the dump channel was tested on APC-C7 (same as LIVE/DEAD NIR) for version 6 and 7 of the panel. Therefore, CD40$^+$ and CD14$^-$ cells were selected as our "classical" B cells (Fig. 1).

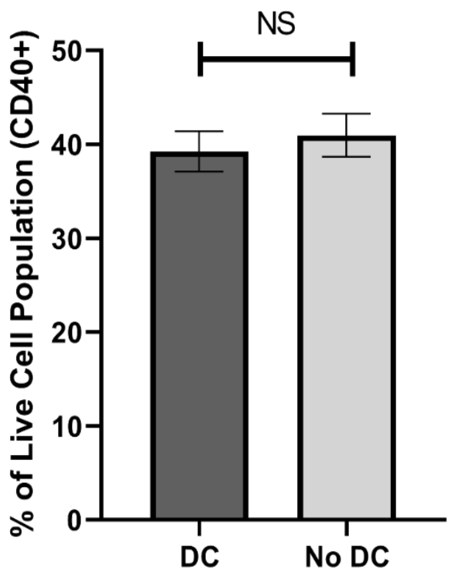


**Online Figure 8** Comparison of the percentage of live gated cells that are CD40$^+$ with and without the dump channel. There was no significant difference between the percentage of CD40$^+$ live cells with or without the dump channel. This was done in triplicate.

Additionally, if intracellular staining is desired, we would recommend custom conjugation of the CD14 and CD71 antibodies to some fluorochromes in the violet laser's line of the instrument. If there is no desire to study monocytes with the B cells, it is also possible to move the CD14 antibody into APC-Cy7 (same as

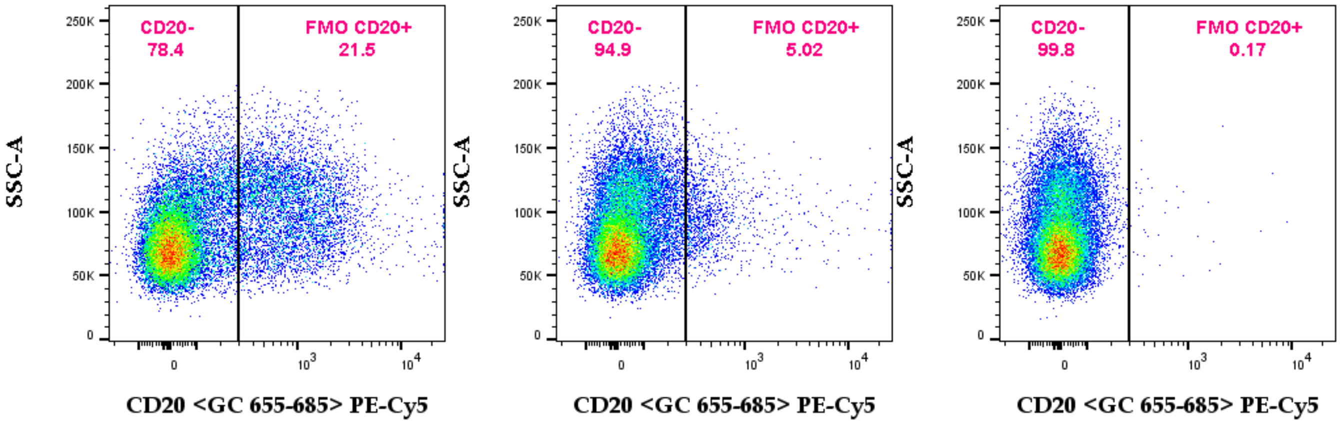


**Online Figure 9** Comparison of different staining strategies for the resolution of CD20 Left-Right: CD20 antibody stained first and followed by staining with all other antibodies; All antibodies in a single mix; CD20 antibody stained after staining with other antibodies.

LIVE/DEAD NIR). It is important to remember that the CD20 antibody must be added first, and only after labelling with the CD20 should one proceed with labelling using the rest of the antibodies in this panel. CD20 did not have good resolution on PE-Cy5 (GC 655-685 detector, version 1-4, re-tested in version 6). Troubleshooting lead to labelling with the CD20 antibody before, at the same time and after the other antibodies (Online Figure 9). CD20 had a radically different staining profile if the CD20 antibody was used to stain the cells before the other antibodies and resembled the single colour staining profile (Online Figure 10). However, changes in the panel forced the change of the CD20 antibody on to PE (GE 575-590 detector, version 5 and 6). However, the CD20 antibody did not perform well on PE as the spectral spread bled into PE-Texas Red (GD 600-620 detector) (Online Figure 7). Thus, it was moved to PE-Cy 7 (version 7) as outlined earlier.

Furthermore, the panel can easily be adapted by adding other markers such as Ki69 and FOXP3 if desired, as these two markers are useful to further inform on the level of activation and differentiation of cattle B cells.

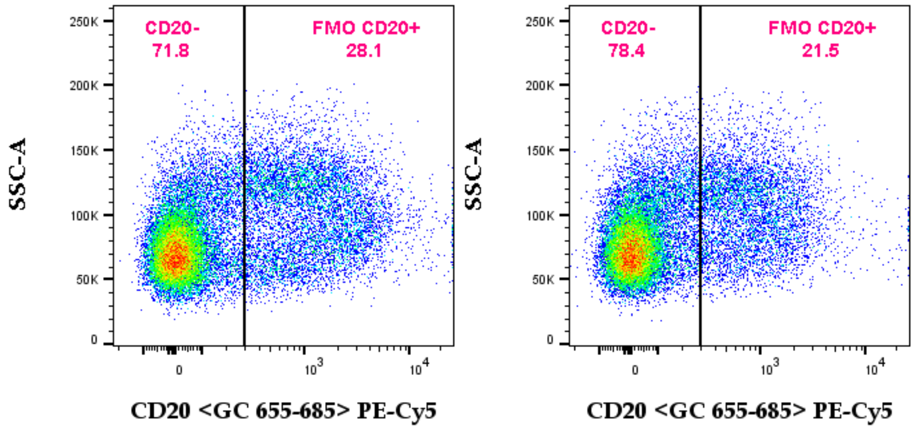


**Online Figure 10** Comparison of CD20 antibody single colour staining (Left) and CD20 antibody stained before all other antibodies (Right).

*FMO controls*

The fluorescence minus one (FMO) control was used to set gates and determine amount of spectral spread between channels that are known to have some spread between them (Online Figure 11 and 12). This indicated that there was some spectral spread between the channels; however, the FMO controls allowed us to set the

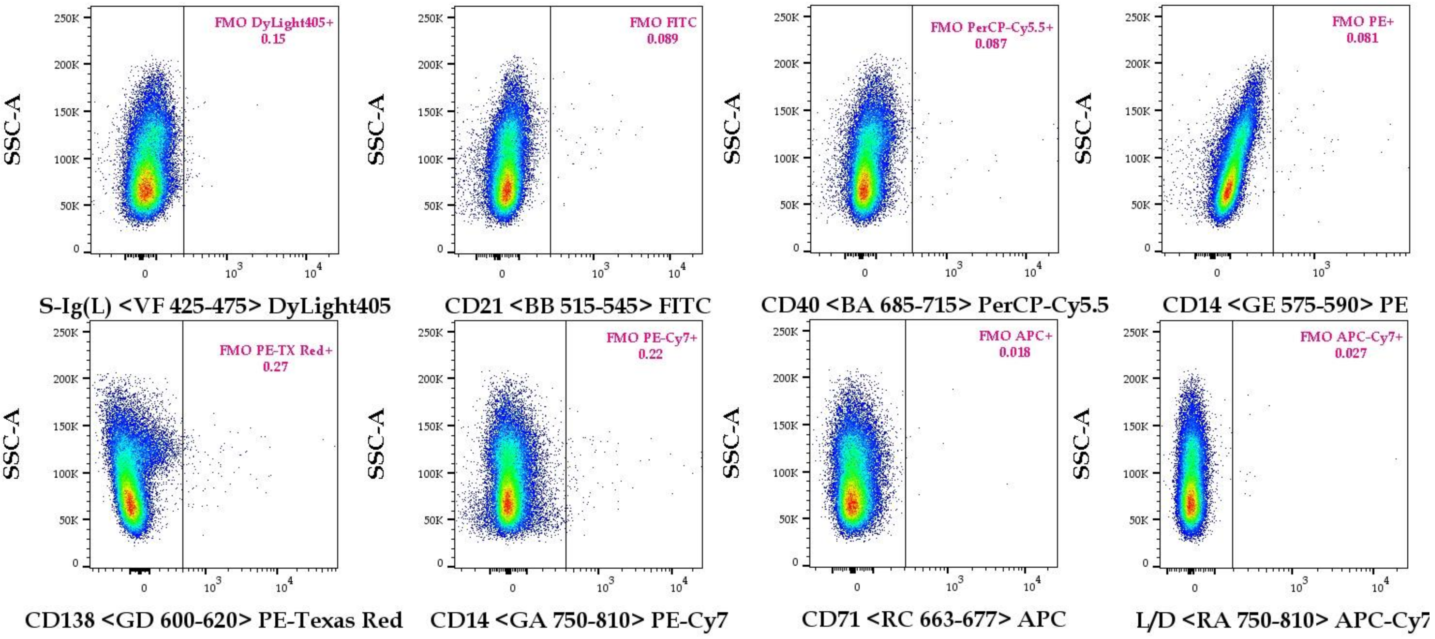


**Online Figure 11** Appropriate fluorescence intensity gating, using the fluorescence minus one control for all fluorochromes in this panel against the side scatter.

proper gates for the populations of interest. Importantly in version 7 of the panel, we did not lose any resolution of the targeted subsets.

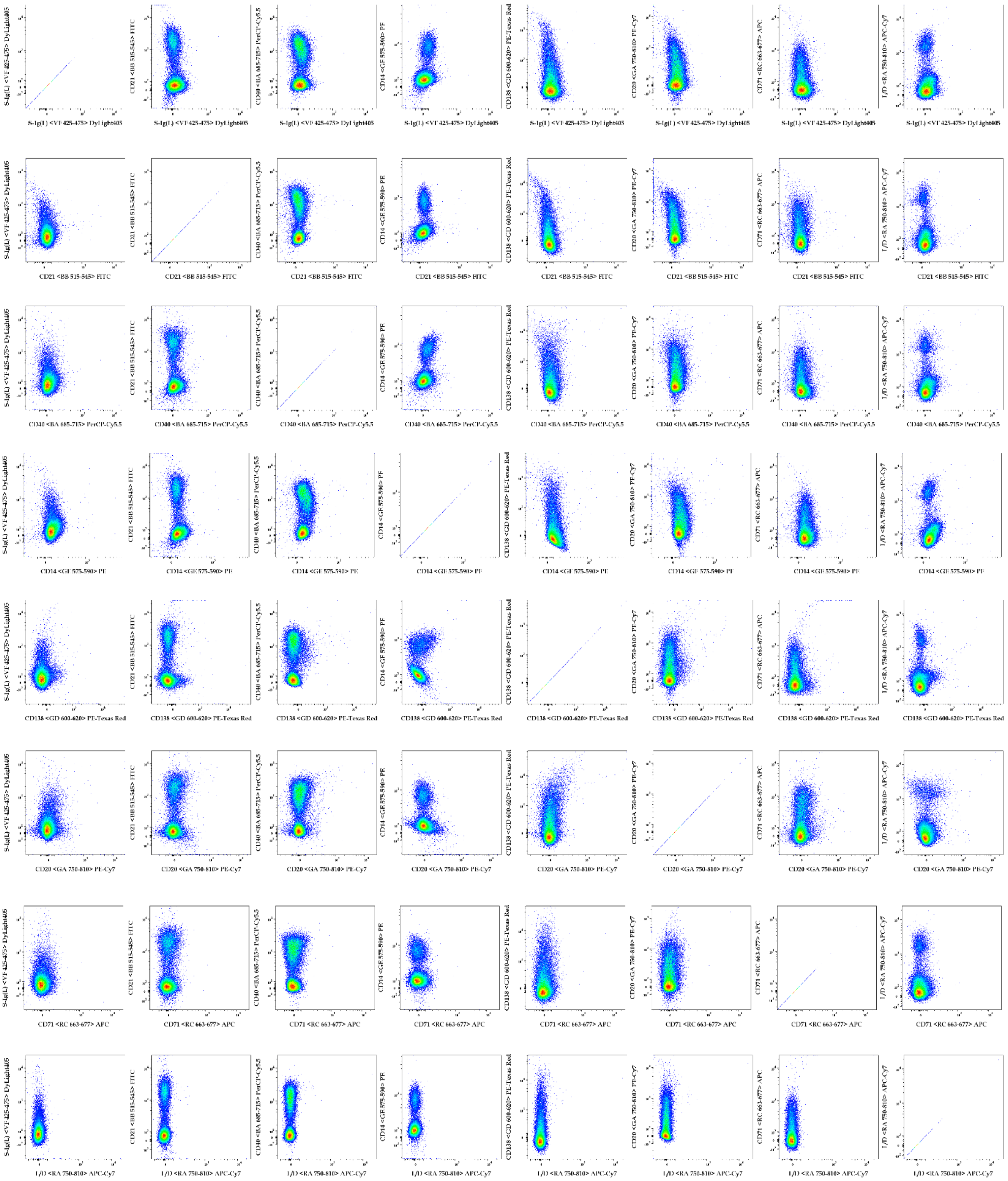

**Online Figure 12** Fluorescence minus one control matrix comparing all fluorochromes in this panel.

*Antibody titration*

The Ab titration was set up by adding 4ul of the 1mg/ml in-house conjugated antibody to 250ul of staining buffer. The antibodies were then titrated in a 2-fold dilution series (1:62.5 1:125, 1:250, 1:500, 1:1000, 1:2000, 1:4000, 1:8000 and an unstained sample) to determine the optimal concentration for each in-house conjugated antibody. The optimal concentration was selected as the lowest concentration with the highest resolution of the population of interest with the least effect on the negative population (Online Figure 13). It is important to note that each new conjugate generated using in-house conjugation kits requires titration prior to use in this panel. Thus, these dilutions are just representative of what was used in the final version of the panel.

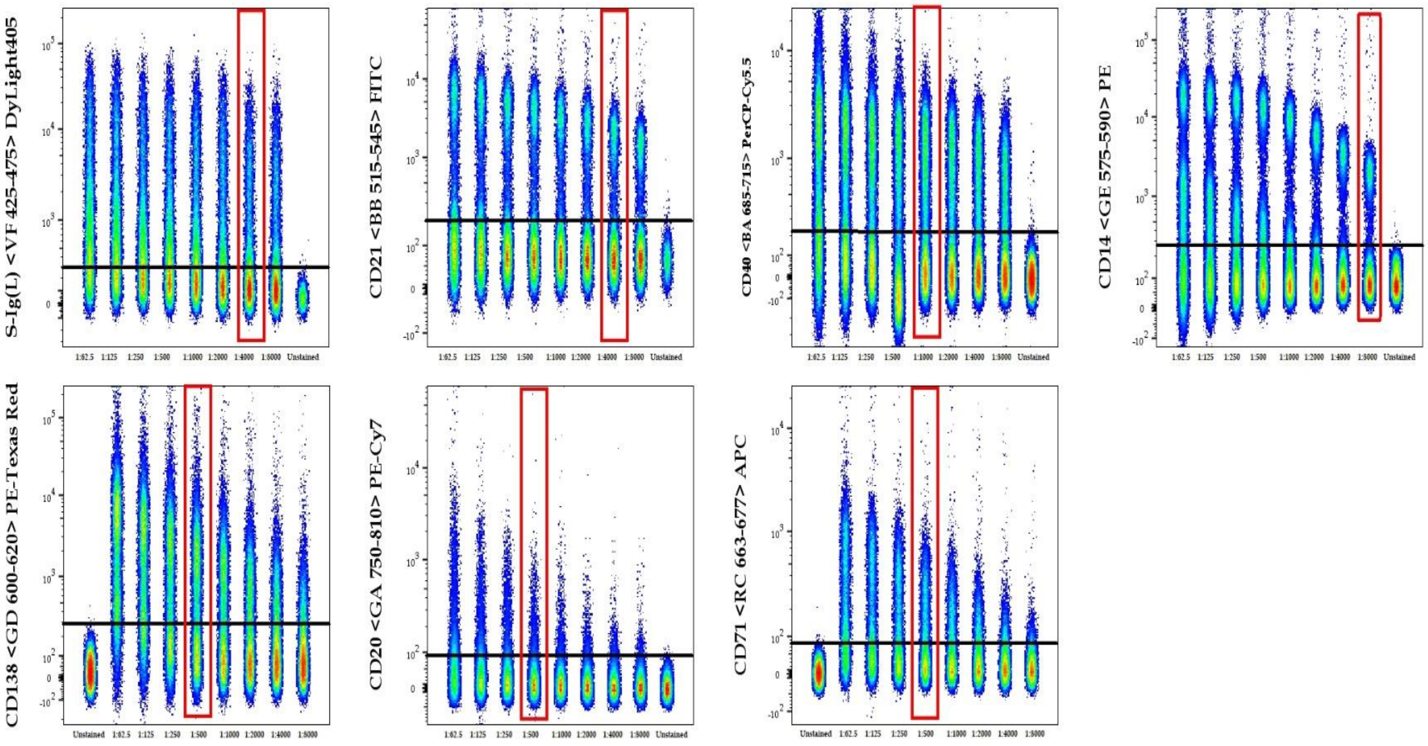


**Online Figure 13** Antibody titrations. Seven 2-fold dilutions were performed. Each concatenated file included an unstained sample to compare the shift of the negative population due to background staining (indicated by the black line). The red box indicated the optimal dilution used in this panel of the specific in-house conjugation.

*Gating strategy*

To differentiate between the different B cell subsets, we first differentiated the subsets on their staining profiles for S-Ig(L) and CD21. We identified the four quadrants as follows; 1: S-Ig(L)$^{+}$CD21$^{-}$, 2: S-Ig(L)$^{+}$CD21$^{+}$, 3: S-Ig(L)$^{-}$CD21$^{+}$ and 4: S-Ig(L)$^{-}$CD21$^{-}$ (Fig. 1). There after each of the quadrants were sub-divided based on the surface staining for CD20 and CD71; 5: CD71$^{+}$CD20$^{-}$, 6: CD71$^{+}$CD20$^{+}$ and 7: CD71$^{-}$ (Fig. 1). These resulting 12 different B cell sub-populations were then further differentiated using CD138, as CD138$^{+}$ or CD138$^{-}$ (Fig. 1). Thus, the panel allows for the classification of 24 unique subsets of cattle B cells. Depending on the subsets of interest, the inverse of this can also be done, in other words the initial differentiation can be done by plotting CD20 against CD71 (5-7) followed by S-Ig(L) against CD21 (1-4), and again lastly looking at the CD138 differentiation of these subsets (Fig. 1).

**Staining Protocol**

**Commercial materials:**

Corning™ Falcon™ 15mL Conical Centrifuge Tubes (VWR International, Cat No. 734-0452)

Horse serum 500 ml (Sigma, H1138)

PBSa (In-House supply)

Polystyrene 96 well conical ´V´ bottom plate (Elkay Laboratory Products (UK) Ltd., Cat No. MICR-TPV)

Autotube 1.1ml tubes (Elkay Laboratory Products (UK) Ltd., Cat No. 000-MICR-120)

**Cell seeding procedure:**

1. Add cells to 10ml of 10% horse serum and PBS in 15ml Falcon tube for 3 min and spin at 300×g for 15 min at 4°C.
2. Re-suspended cells in PBS and aliquot 100µl to each well (96 V-well plate) so that each well contains $1\times10^6$ cell/well.
3. Centrifuge plate at 300×g for 5 min at 4°C (for all centrifugation steps below).

**Staining:**

4. Add 100µl of conjugated CD20 (PE-Cy7) at 1:500 to cells and, resuspend by gently pipetting.
5. Incubate 15 min at RT in dark.
6. Centrifuge and wash ×2 with 200µl of PBS.
7. Add 100µl of conjugated antibody mix to each sample well, resuspend by gently pipetting.
8. Incubate 15 min at RT in dark.
9. Centrifuge and wash ×2 with 200µl of cold PBS.
10. Add 100µl of LIVE/DEAD NIR diluted 1:2000 in cold PBS to cells, resuspend by gently pipetting.
11. Incubate 10min at RT in dark.
12. Centrifuge and wash ×2 with cold PBS only.
13. Re-suspend pellet, by gently pipetting 200µl cold PBS and transfer to 1.1ml tubes.

**Literature Cited**

1. Ellebedy AH, Jackson KJL, Kissick HT, Nakaya HI, Davis CW, Roskin KM, McElroy AK, Oshansky CM, Elbein R, Thomas S, Lyon GM, Spiropoulou CF, Mehta AK, Thomas PG, Boyd SD, Ahmed R. Defining antigen-specific plasmablast and memory B cell subsets in human blood after viral infection or vaccination. Nat. Immunol. 2016;17:1226–1234. Available at: http://www.nature.com/articles/ni.3533.

2. Cossarizza A, Chang H-D, Radbruch A, Acs A, Adam D, Adam-Klages S, Agace WW, Aghaeepour N, Akdis M, Allez M, Almeida LN, Alvisi G, Anderson G, Andrä I, Annunziato F, Anselmo A, Bacher P, Baldari CT, Bari S, Barnaba V, Barros-Martins J, Battistini L, Bauer W, Baumgart S, Baumgarth N, Baumjohann D, Baying B, Bebawy M, Becher B, Beisker W, Benes V, Beyaert R, Blanco A, Boardman DA, Bogdan C, Borger JG, Borsellino G, Boulais PE, Bradford JA, Brenner D, Brinkman RR, Brooks AES, Busch DH, Büscher M, Bushnell TP, Calzetti F, Cameron G, Cammarata I, Cao X, Cardell SL, Casola S, Cassatella MA, Cavani A, Celada A, Chatenoud L, Chattopadhyay PK, Chow S, Christakou E, Čičin-Šain L, Clerici M, Colombo FS, Cook L, Cooke A, Cooper AM, Corbett AJ, Cosma A, Cosmi L, Coulie PG, Cumano A, Cvetkovic L, Dang VD, Dang-Heine C, Davey MS, Davies D, De Biasi S, Del Zotto G, Dela Cruz GV, Delacher M, Della Bella S, Dellabona P, Deniz G, Dessing M, Di Santo JP, Diefenbach A, Dieli F, Dolf A, Dörner T, Dress RJ, Dudziak D, Dustin M, Dutertre C, Ebner F, Eckle SBG, Edinger M, Eede P, Ehrhardt GRA, Eich M, Engel P, et al. Guidelines for the use of flow cytometry and cell sorting in immunological studies (second edition). Eur. J. Immunol. 2019;49:1457–1973. Available at: http://doi.wiley.com/10.1002/eji.201646632.

3. Baker D, Marta M, Pryce G, Giovannoni G, Schmierer K. Memory B Cells are Major Targets for Effective Immunotherapy in Relapsing Multiple Sclerosis. EBioMedicine 2017;16:41–50. Available at: http://dx.doi.org/10.1016/j.ebiom.2017.01.042.

4. Weisel F, Shlomchik M. Memory B Cells of Mice and Humans. Annu. Rev. Immunol. 2017;35:255–284. Available at: https://www.annualreviews.org/doi/10.1146/annurev-immunol-041015-055531.

5. Mwangi W, Maccari G, Hope JC, Entrican G, Hammond JA. The UK Veterinary Immunological Toolbox Website: promoting vaccine research by facilitating communication and removing reagent barriers. Immunology 2020;161:25–27. Available at: http://doi.wiley.com/10.1111/imm.13227.
